# Conservative method for attenuation of Vertical Electrooculogram based on local suppression of artifact templates from ongoing EEG

**DOI:** 10.1101/268763

**Authors:** Dimitri M. Abramov, Carlos Alberto Mourão, Paulo Ricardo Galhanone, Vladimir V. Lazarev

## Abstract

**Background:** Eye movement during blinking can be a significant artifact in ERP analysis (mainly if ERP is blink-locked). Blinks produce a large positive potential in the vertical electrooculogram (VEOG), spreading towards posterior direction. Two methods are the most frequently used to suppress VEOGs: linear regression to subtract the VEOG signal from each EEG channel and Independent Component Analysis (ICA). However, both lose some EEG information.

**Methods:** The present algorithm (1) statistically identifies the time position of VEOGs in the frontopolar channels; (2) performs EEG averaging for each channel, which results in ‘blink templates’; (3) subtracts each template from the respective EEG at each VEOG position, only when the linear correlation index between the template and the segment is greater than a threshold L. The signals from twenty subjects were acquired using a behavioral test and were treated using FilterBlink for subsequent ERP analysis. A model was designed to test the method for each subject using twenty copies of the EEG signal from the mid-central channel of the subject (which has almost no VEOG) representing each one of the 20 EEG channels and their respective blink templates. At the same 200 equidistant time points (marks), a signal (2.5 sinusoidal cycles at 1050 ms to emulate an ERP) was mixed with each model channel, along with the respective blink template of that channel, between 500 to 1200 ms after each mark.

**Results:** According to the model, VEOGs interfered with both ERPs and the ongoing EEG mainly on the anterior medial leads, and no significant effect was observed on the mid-central channel (Cz). FilterBlink recovered approximately 90% (at Fp1) to 98% (Fz) of the original ERP and EEG signals to L of 0.1. In the analysis of real signals, the method reduced drastically the VEOG effect on the EEG after ERP and blink-artifact averaging.

**Conclusion:** The method is very simple and effective for VEOG attenuation without significant distortion of the EEG signal and embedded ERPs in all channels.

## INTRODUCTION

Vertical eye movement during blinking produces a positive and symmetrical potential on the electroencephalogram signal, the vertical electrooculogram (VEOG), with the greatest amplitude at frontopolar (Fp) sites (where it is easily observable), which decays by conduction along the scalp surface [1]. In ERP studies, during signal averaging, such a large potential can be a significant artifact that would interfere with waveforms, mainly at anterior scalp sites. If the blink is a reflexive and systematic response to testing design (such as attentional tests that promote expectancy), and it is time locked to the studied events, even the low amplitude VEOG at posterior channels could reveal a significant effect upon the event-related EEG potentials [2, 3].

Parallel to the development of human neurophysiology, methods for minimizing the VEOG effect on neural signals have been implemented. Digital signal recording has allowed several off-line signal processing strategies to suppress VEOG effects. A usual method uses the electrooculogram signal (EOG-S) recorded from the ocular channel (around the eyes) as a template to extract the VEOG after a linear regression between the latter and each channel, and its corresponding correlation coefficient reduces EOG-S amplitude, which is subtracted from the EEG signal [4, 5, 6]. However, this method corrupts the neural information as the EOG-S also contains EEG signal, which is also subtracted by the algorithm [7].

Another method offers more effective and reliable results, although it is computationally more intensive: correction using an Independent Component Analysis (ICA) [7, 8]. The algorithm for ICA-based EEG correction needs an external classifier to choose which components refer to the EOG-S [7, 8, 9, 10]. However, ICA has a serious issue with regards to artifact removal [11]: as there are fewer channels on the scalp, there is more neural information in the components than in the artifact signal. Thus, even a conservative method such as ICA could corrupt the ERPs of interest if the EEG setup has twenty channels (conventional 10-20 system), for example.

To develop an automatic algorithm for the suppression of VEOG which is suitable for both the ERP analysis (which works on time domain with minimum EEG corruption) and for datasets with few channels, we have implemented a simpler approach that attempts to suppress only the VEOG artifacts from the EEG. The method is based on the subtraction of the “blink template” (processed from the frontopolar channels) from the original EEG in each channel provided that the correlation coefficient between them exceeds a predefined threshold.

## Methods

### Experimental Procedures and ERP acquisition

Twenty typically developing boys (9-13 y.o.) were submitted to the Attention Network Test [12], a forced choice protocol with two alternatives for evaluation of target orientation (a yellow fish facing right or left), chosen by pressing the respective keyboard arrows. The target position in the visual field is indicated by different kinds of cues. Reaction time, response accuracy, and long latency ERPs, recorded at the parietal and frontal sites, were studied. The ANT is divided into trials with 1650ms of stimulus between cue and target onsets, separated by gaps randomly varying between 1000 and 2000 ms, after the behavioral response. In each trial, the subject was asked to fix their glance at a central cross, and immediately press with his finger the left or right arrow key of the keyboard, according to the target horizontal orientation. (a yellow fish)., Each subject performed nine blocks (a training block plus eight test blocks), with 24 trials each (for all possible cues and target conditions).

The setup Neurofax^®^ 1200 (Nihon Kohden) with 20 channels (10-20 montage system) was used, with Ag-AgCl electrodes over the scalp, referenced at A1-A2. The impedances were maintained below 10 μΩ Target stimuli were synchronized to the EEG by a digital trigger from the computer that displayed the ANT screens and recorded the motor responses. The neural signal was digitalized with a sampling rate of 1000Hz, and 24-bit resolution. High- and low-pass filters were, respectively, 0.5 Hz and 150 Hz, with a FFT notch filter at 60Hz. These filters were applied after acquisition of signals (offline). The remaining relevant noise (e.g., muscular artifacts) was eliminated by manual inspection of the EEG signal prior to averaging and after FilterBlink application.

### The FilterBlink algorithm

The present method suppresses VEOG artifact by subtracting its “template” (i.e. the grandaverage of all detected VEOGs from the respective channel) from an EEG segment when it shows sufficient correlation with the template. It is obtained for each channel by averaging the VEOGs, which are triggered at the frontopolar channels (Fp1 and Fp2). The time positions where the blinks occur are detected when the signal amplitude exceeds a threshold given by the mean, *x̄*, plus *n* standard deviations, σ, of the EEG signal modulus in the Fp1 and Fp2 channels. Therefore, from these EEG signals, of size M, the vector of the VEOG positions, *P*, of size *N*, was extracted for EEG amplitude values greater than *n*σ + *x* (here, we have arbitrarily chosen *n* = 1.5). All points, *pi*, between *i* and *i* + *f*/10 (where *f* is the sampling frequency of the digital signal) have been discarded as they belong to the same VEOG with period always greater than 100ms. Hence, the wave points of the vector *P´* were selected as:

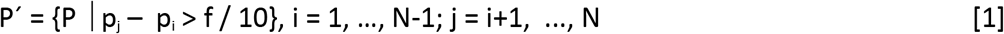

Two vectors of wave points, *P’*, are identified (for Fp1 and Fp2). The rate of their lengths is a measure of confidence. If it lay in an arbitrary interval, *d* (here, 0.9 < d < 1.1), the detection of VEOGs would be regarded as valid. Otherwise, FilterBlink would be aborted.

The shorter P’ vector is assumed as the indexing vector of the central positions of the VEOGs, which is applied to all channels. Hence, the EEG signal for each one of the 20 channels, X, is averaged inside an epoch p’_i_ – t and p’_i_ + t, yielding the VEOG templates, Θ (vector with amplitude values at the time, for each channel, j, from the channel set, X), using the following averaging algorithm:

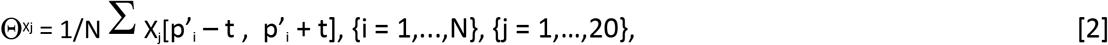

where *t* = 350ms. The linear correlation coefficients, *r*, are obtained when the respective VEOG template and the EEG epoch from an EEG channel *X*_*j*_ are correlated using Pearson’s method. Whenever *r* > *L*, the template Θ from the EEG channel *X*_*j*_ is subtracted from the respective epoch:

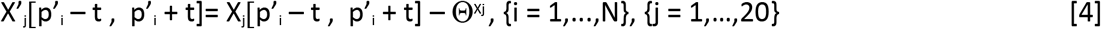

The adopted values for *L* are 0.1, 0.2, 0.3, 0.4, 0.6, and 0.8. See figure 01 for a diagram explaining the FilterBlink operating algorithm.

**Figure 01.**
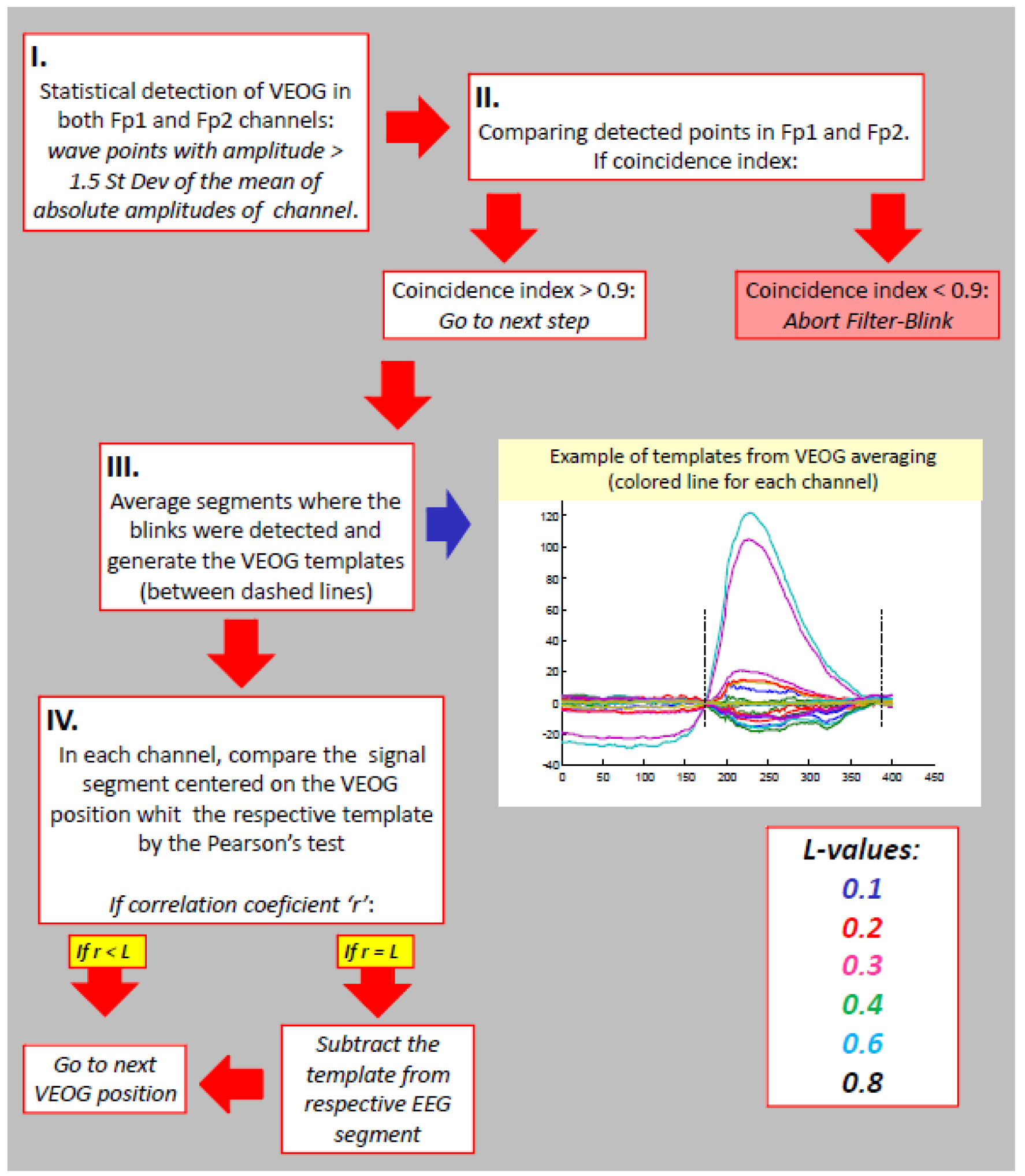
Diagram of the FilterBlink operating algorithm.

### The EEG Model

To verify its efficacy, the present method was tested using an EEG model with embedded artificial event related potentials (ERPs) with predetermined parameters. A model was performed for each subject using 20 copies of his Cz signal, where the blink artifact is relatively irrelevant. These 20 copies are the base for the modeled channels, equivalent to those from the system. A trigger vector with 200 markers spaced 4400 ms apart was generated. A sinusoidal wave with 2.5 cycles (0: 5πrad), −5 to 5 μV of amplitude, and 1150 ms long was inserted 100ms after every marker onset, in all model channels, representing the ERP. The real VEOG templates from the respective subject, per each channel, were added to their respective modeled channels after each marker, with their position randomly ranging from 500 to 1200 ms (the frequency of random values was normally distributed). The amplitude of the VEOG templates was stochastically enlarged up to 1.4 times. See figure 02.

**Figure 02.**
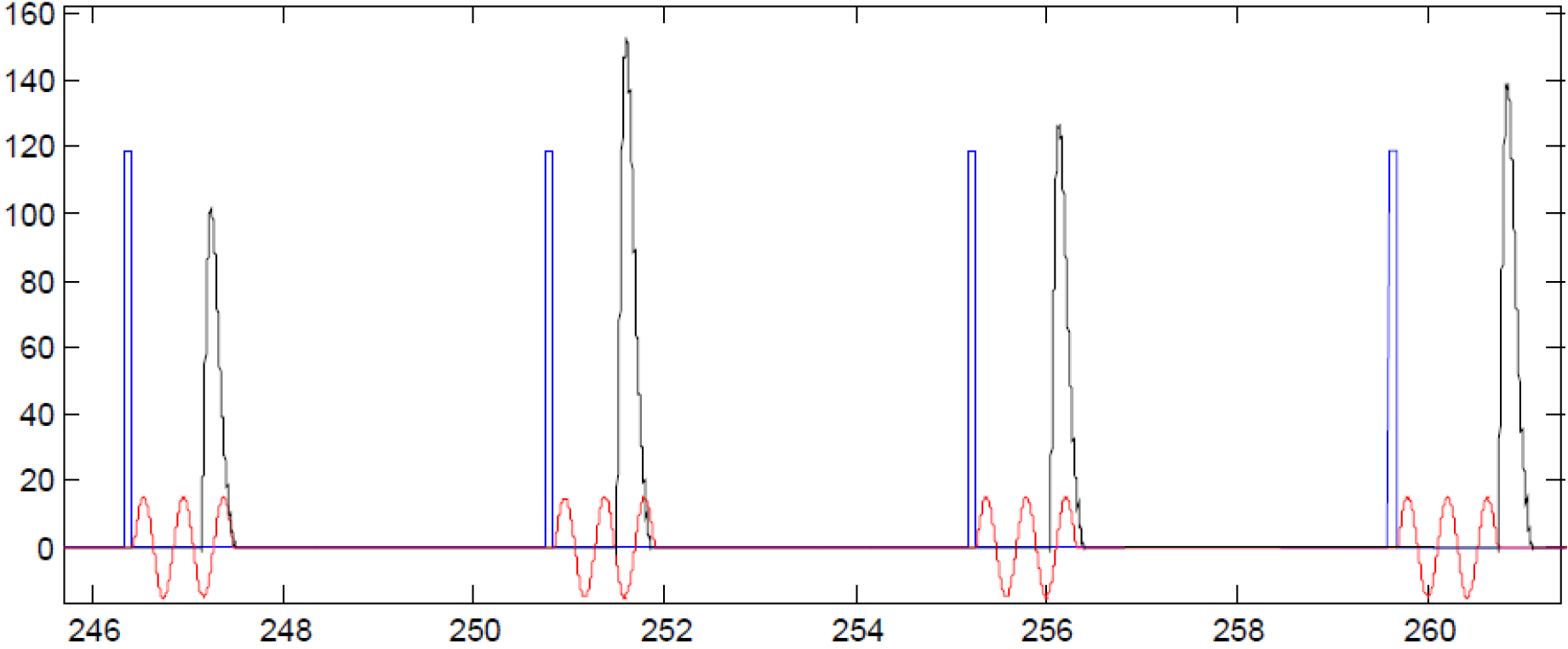
Schematic illustration of the generation of the modeled EEG signal.

### Statistical analysis

Pearson’s method was used to test correlations between modeled EEG signal without VEOG inclusion (only the artificial ERP wave was embedded), EEG signal with included VEOG artifact, and modeled EEG signal after FilterBlink application, for each L value. The correlation coefficients formed a dataset for statistical comparisons between these three conditions using the Wilcoxon Signed Rank Test for paired variables.

The results of FilterBlink application in the real EEG were qualitatively described for the ongoing EEG at frontopolar channels, as well as the effects of FilterBlink on ERP waves and averaged blink artifacts.

## Results

### Quantitative Analysis of Model Results

The present method apparently eliminated most of the blink artifacts in the modeled EEG signal of the Fp1 channel (figure 3A).

**Figure 03.**
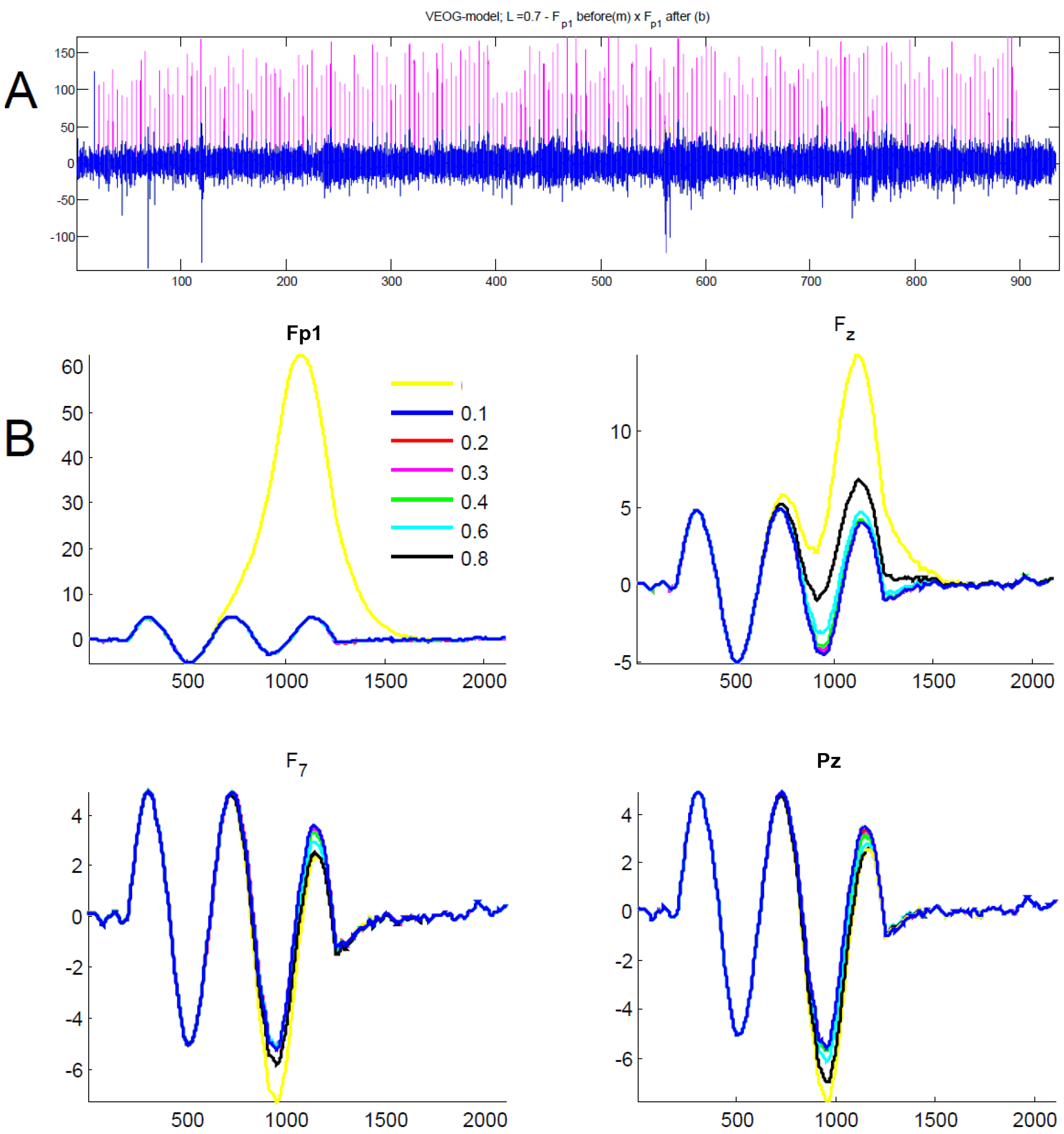
Application of FilterBlink on the EEG modeled signal. A. Illustrative effect on the Fp1 modeled channel, prior to (magenta) and after FilterBlink application. B. After signal averaging is locked to trigger vector, the effect of each L value on FilterBlink application at Fp1, Fz, F7, and Pz modeled channels, showing best ERP (sinusoidal) recovering for L = 0.1 (see color legend). Yellow = without FilterBlink.

Paired comparisons of mean correlation indexes of modeled signals with and without VEOG inclusion, prior to and after FilterBlink application (figigure 4A, red line), revealed that the modeled signal at Cz channel had no prominent alteration (p > 0.05, n = 20) and FilterBlink did not alter this correlation for all *L* values (figure 4B, red line). However, in the other channels, the inclusion of the VEOG templates yielded low correlation coefficients between the signals prior to and after adding VEOG artifacts (figure 4A). And FilterBlink application recovered the similarity with the signals prior to artifact inclusion, when coefficients turn from 0.985 to 0.995, for L = 0.1 (p < 0.001, Wilcoxon Rank Test, for Fz, Pz and Oz, figure 4A). The higher L values yielded a progressive reduction of similarity between the modeled signal prior to artifact inclusion and after FilterBlink application, with a logarithm fashion (p < 0.001 comparing waves for L= 0.1 and L > 0.6, Wilcoxon Rank Test, for Fz, Pz, and Oz, figure 4B).

**Figure 04.**
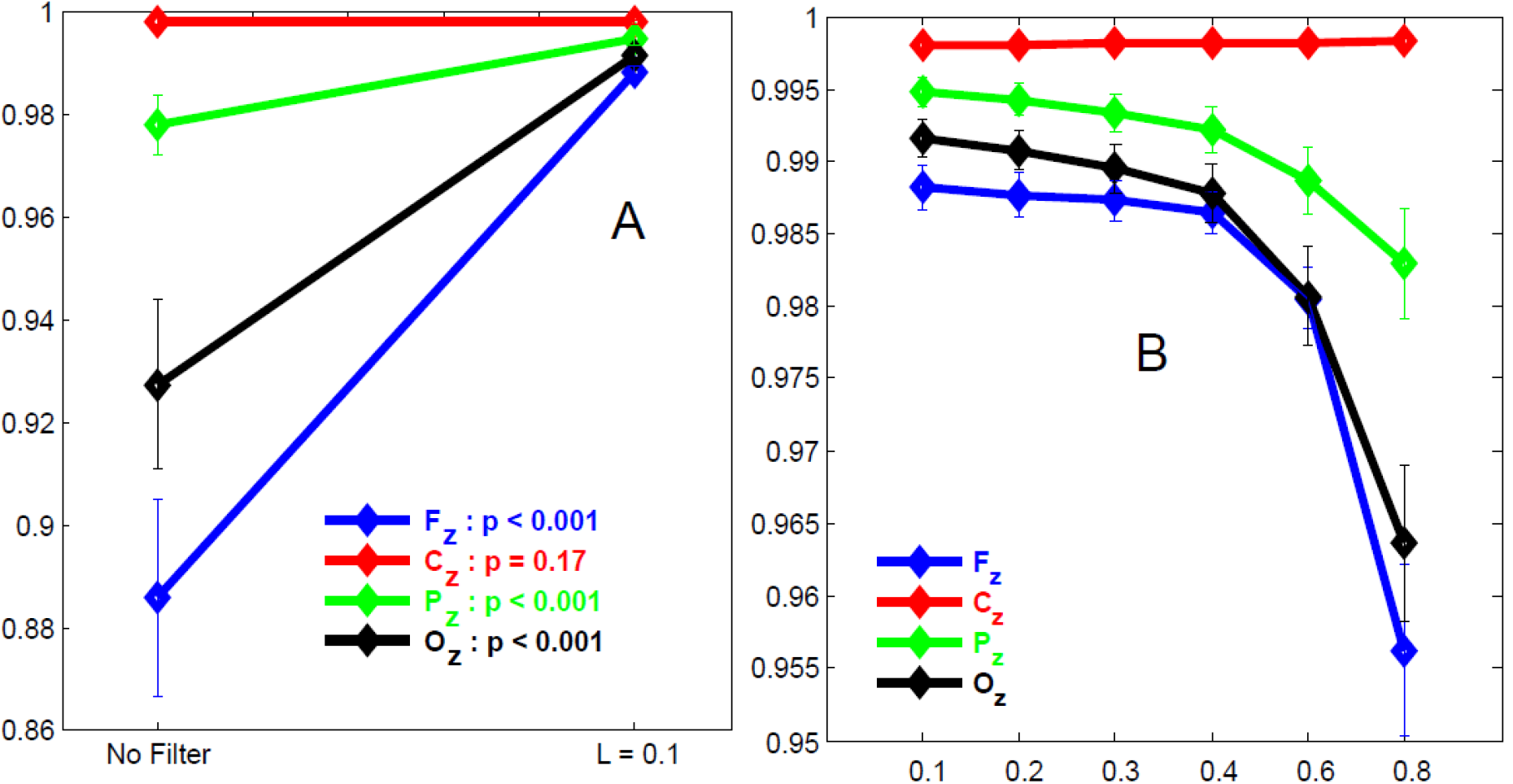
Analysis of the FilterBlink effect on the correlation index between modeled ongoing EEG without VEOGs and modeled ongoing EEG with VEOGs. A. Correlation index prior to and after FilterBlink application (L=0.1). B. Correlation index for different L values (sample mean ± std. dev.).

As observed for the ongoing modeled EEG signal, the sinusoidal ERPs were not affected either by artifact inclusion or by the application of FilterBlink at Cz (figure 5A, red line). However, FilterBlink recovered the sinusoidal wave overlaid on the blink and preserved the remaining artificial ERPs (before 500ms in the epochs). The ERPs prior to (figure 3B, yellow line, for channels Fp1, Fz, F7, and Pz) and after FilterBlink application (colored lines for each L value, figure 3B) were substantially different despite the fact that signal distortion by VEOGs seems to be more relevant in Fz and frontopolar channels.

**Figure 05.**
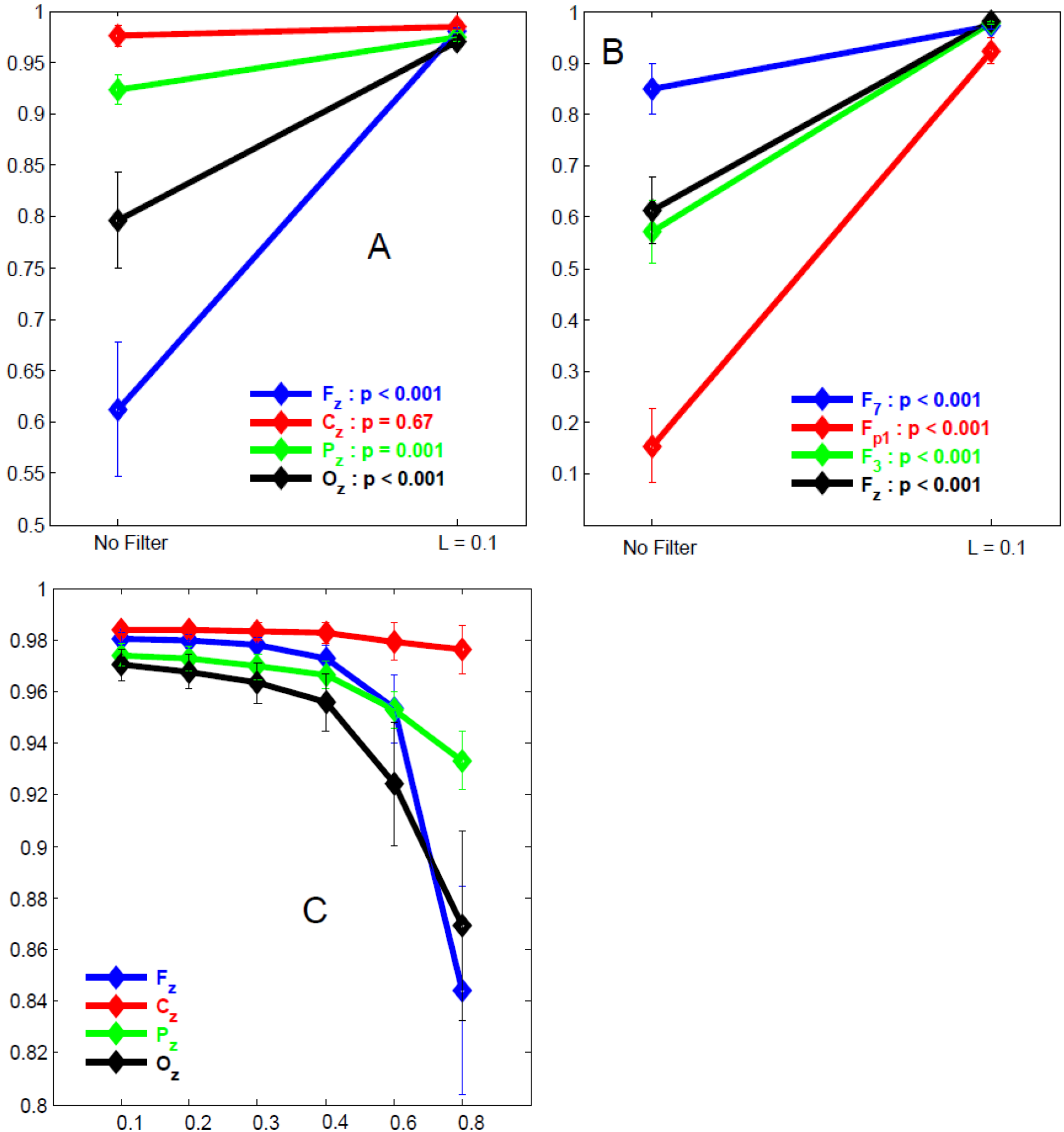
Analysis of the FilterBlink effect on the correlation index between modeled ERPs without VEOGs and modeled ERPs with VEOGs in the ongoing EEG. A. Correlation index prior to and after FilterBlink application (L=0.1). B. Correlation index for different L values (sample mean ± std. dev.).

Comparing Pearson`s coefficients of the correlation between artificial ERPs prior to and after FilterBlink, we observed that signal recovery was confirmed, which showed an important change from r = 0.15 to 0.9 in the Fp1 channel, or a more discrete effect, changing from r = 0.92 to 0.97 in the Pz channel, (see figure 5A and B, p < 0.001 comparing Pearson`s coefficients prior to and after FilterBlink application for L=0.1). The effect of FilterBlink is more substantial on anterior channels (see histograms of the coefficient differences prior to and after applying FilterBlink for all modeled channels in figure 6, showing the magnitude of the FilterBlink effect on whole scalp signals). Similar to the ongoing modeled signal, the increase in L value reduces signal recovery in a logarithmic curve, and Pearson`s coefficients were statistically different from L=0.1 an L > 0.6, for Fz, Pz, and Oz (p < 0.01, figure 5B).

**Figure 06.**
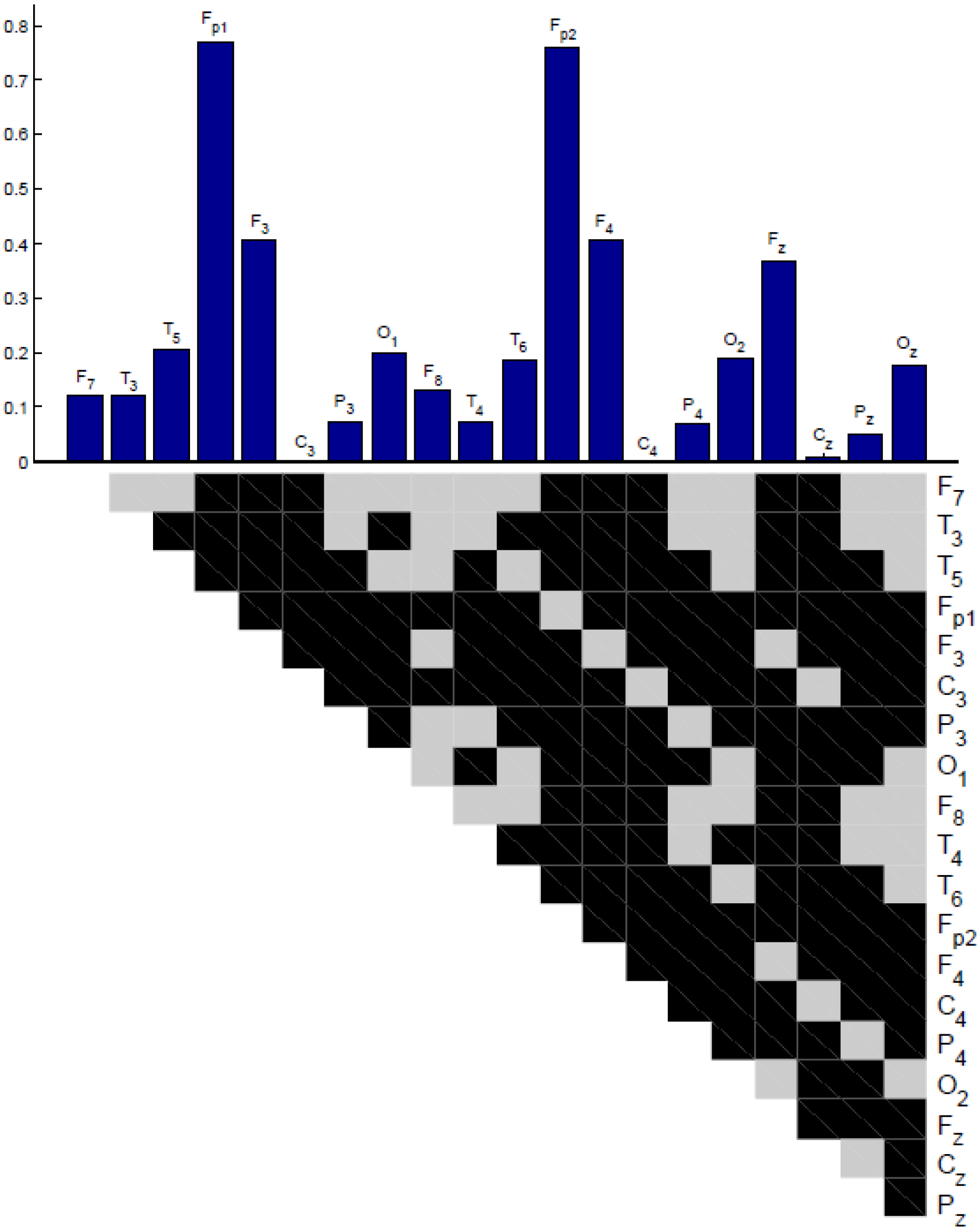
Magnitude of the FilterBlink Effect on modeled ERPs in all channels, i.e., the difference in mean correlation coefficients for ERPs with and without VEOG inclusion in ongoing EEG prior to and after the FilterBlink effect. Black plots represent p < 0.05 for pairwise comparisons of magnitudes between channels (Wilcoxon signed-rank test, without p correction for multiple comparisons).

### Qualitative evaluation of biological data

Artifact suppression occurred in the ongoing EEG recorded during? ANT performance in the Fp1 (presented for 3 subjects in figure 7)prior to (magenta) and after (blue) FilterBlink application (figure 7, for L = 0.1). In a closer view (Fig 7, right column), the EEG background was unaffected by the method while the blink artifacts are subtracted and some non-linear residual oscillation remains.

**Figure 07.**
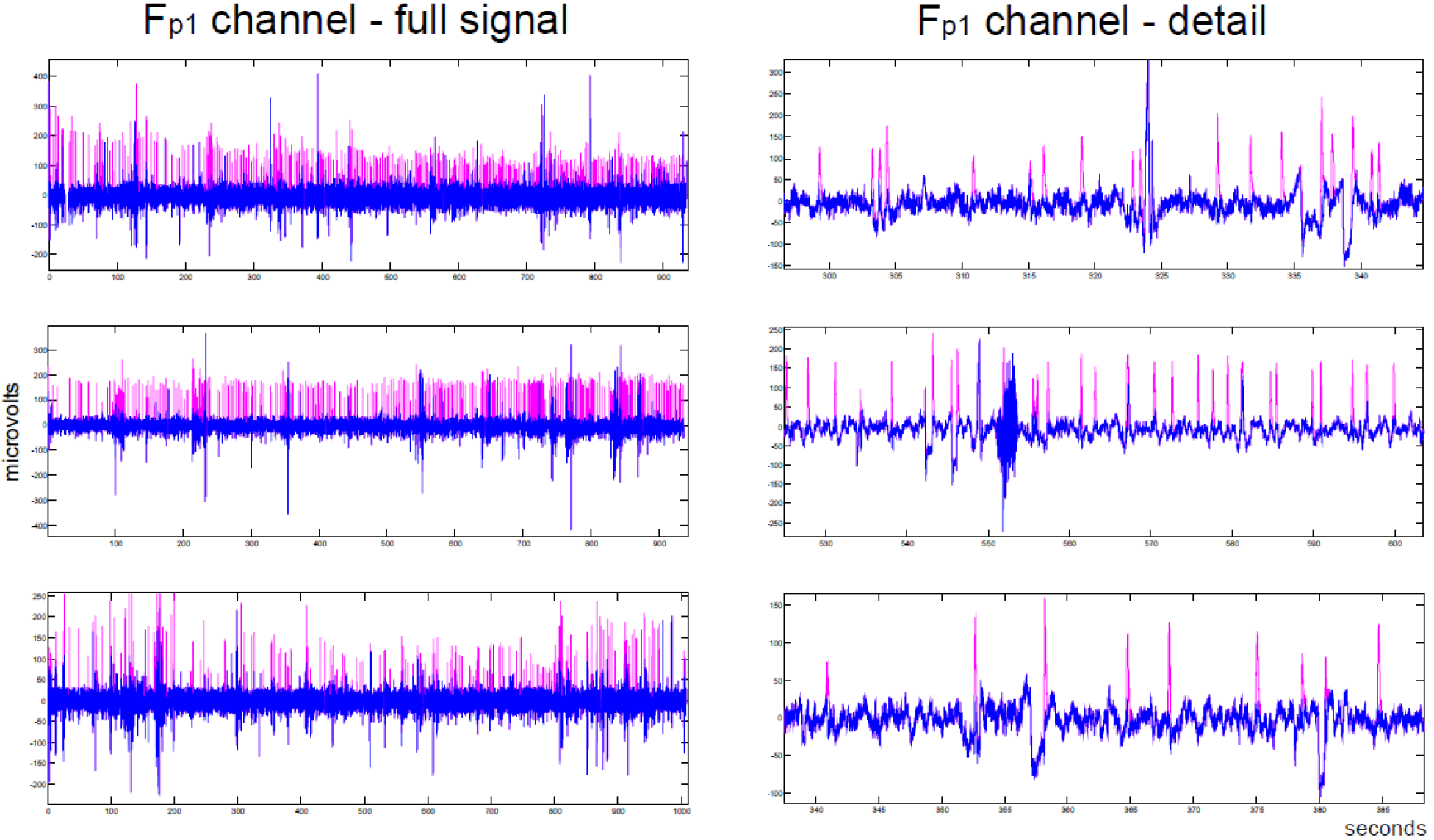
Real ongoing EEG signal prior to (magenta) and after (blue) FilterBlink application in the Fp1 channel, for three subjects with L = 0.1.

After grand averaging, the ERP wave characteristics could be identified in the channels of interest (figure 8: Fp1, F7, Fz, and Pz). Also at the averaged VEOGs (seen in figure 8, yellow wave), the ERPs were restricted to the first 1000 ms in the epochs while the amplitude of the blink artifact increased after that time. The effect of FilterBlink application was more pronounced up to L = 0.4. The averaged VEOG positions (figure 9) confirmed the suppression of blink artifacts up to L < 0.6, where the voltage was near zero line. However, a small positivity/negativity remained after the suppressions in all illustrated channels.

**Figure 08.**
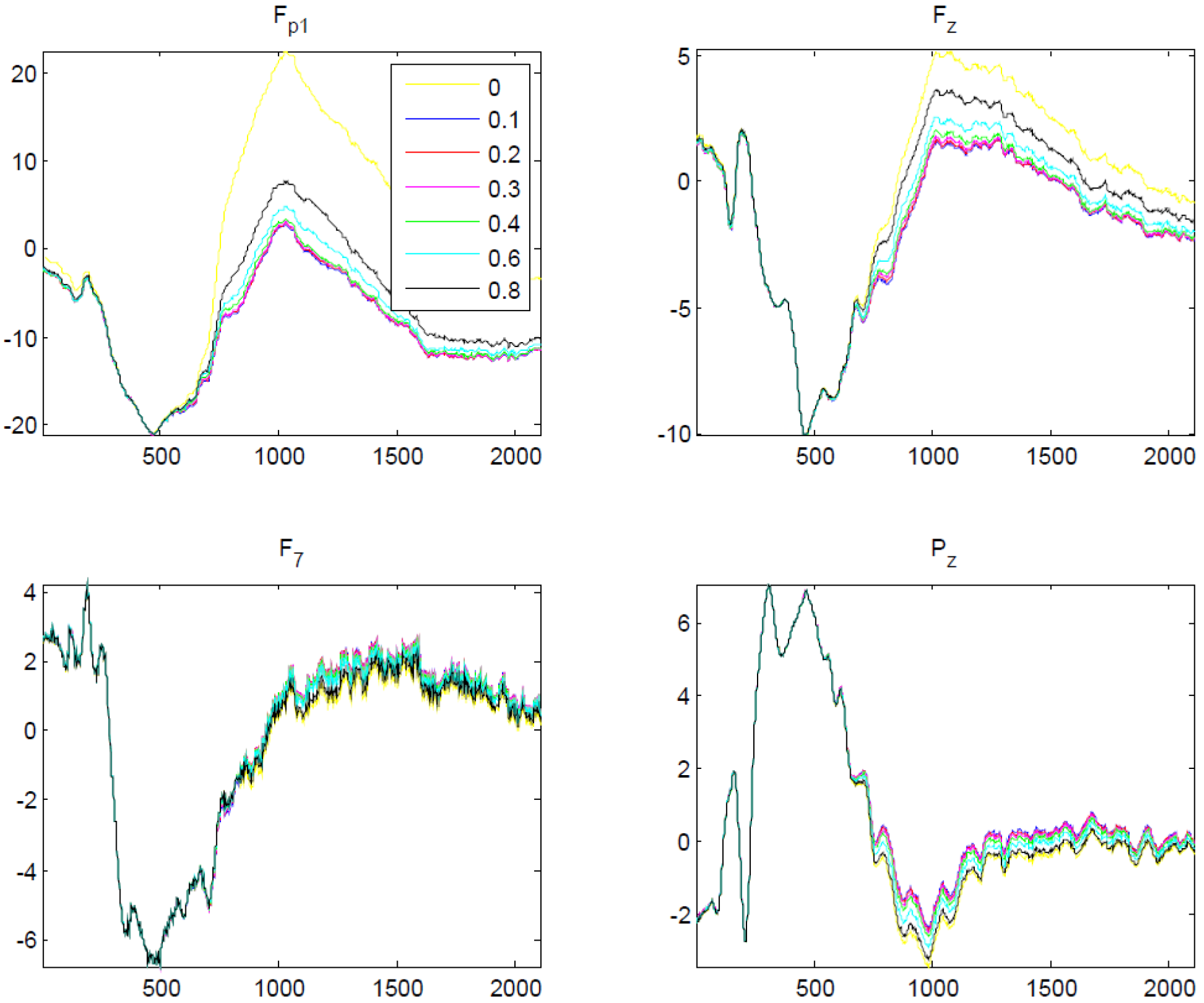
Application of FilterBlink on a real EEG signal and subsequent ERP study with signal promediation. Yellow = without FilterBlink.

## Discussion

Removing artifacts is an important task for ERP analysis when such elements are relatively synchronized to the signals of interest. The main purpose of FilterBlink has been to improve the quality of ERPs which can be degraded by blink artifacts relatively locked to them. The averaging approach is here applied in reverse: VEOG templates obtained with EEG averaging at the time positions where the blink artifacts were detected (from frontopolar channels, with the greatest VEOG amplitude) are subtracted fromthese very time positions; this suppresses only the patterned portion of the artifact, and the solely remaining part is non-linear variations, which naturally disappear with ERP averaging.

The present method is notably simpler than ICA and more conservative than the correction method by regression techniques, since FilterBlink acts locally on the VEOG related EEG signal and does not subtract neural information [4]. For EEG setups with fewer channels, the ICA could corrupt the signal by carrying neural information in the artifact components since there are much more sources than receptors [11]. This does not occur using FilterBlink: it is a process that minimally corrupts the ongoing EEG because the intervention is punctual over a minimal time segment epoch, centered on the positions where the VEOGs were detected, with the same size of the lower portion of the template epoch (only the VEOG deflection, detected between the two minima around the center) used in the process.

**Figure 09.**
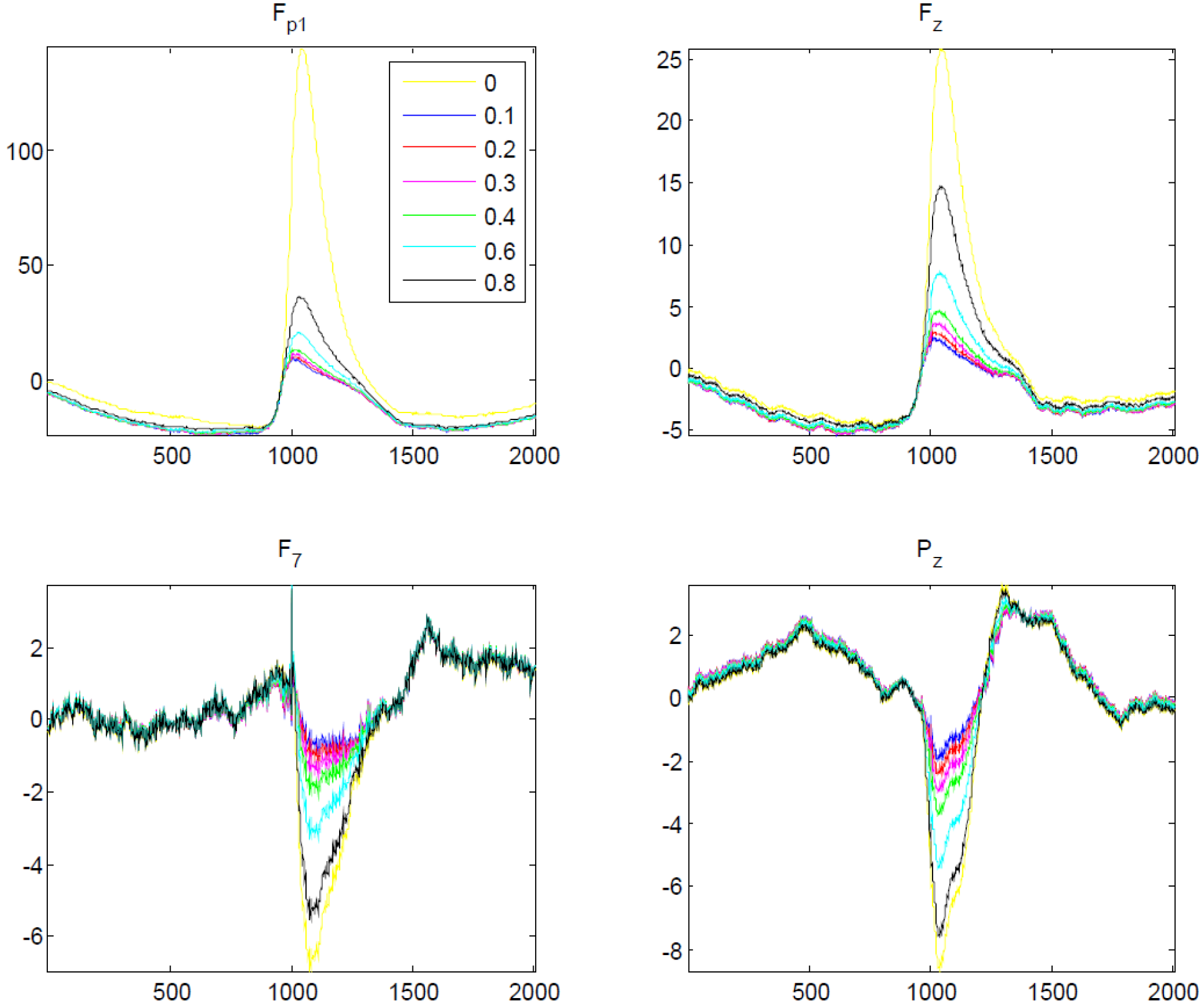
Application of FilterBlink on a real EEG signal and subsequent promediation centered on VEOG detected positions. Yellow = without FilterBlink.

Our method proved to be significantly effective: it is controlled, and it happens in time domain using statistically obtained real elements, without any further signal handling (such as decomposition), and its results could be regarded as predictable and reliable. FilterBlink over the EEG model recovered nearly 98% of the original ongoing signal, and the same result was observed for artificial ERPs, which were partially overlaid by the VEOGs. The model should be regarded as consistent and representative since the biological EEG signal from the mid-central channel of each subject (with the lowest VEOG artifact and with the main brain rhythms [13]) was used, as well as the respective subjects’ VEOG templates for each channel, which are also biological, real elements. The artificial sinusoidal wave for ERP emulation, with low amplitude, was a known pattern for the accurate evaluation of the method after averaging.

As a matter of fact, there are statistical approaches to algorithm processing, obviously because the EEG signal is a complex dataset with chaotic behavior. However, estimates are restricted to the detection of VEOGs, which is performed by marking points on the wave with amplitude greater than *n* standard deviations. If the signal is previously treated to eliminate other artifacts with large amplitude and filtered for long-range oscillations with frequency smaller than 0.5Hz, the error tends to be minimal if an adequate value for *n* is chosen (1.5 standard deviation is usually suitable). The *L* parameter (the threshold of similarity applied to the Pearson’s test coefficient regarding the correlation between the template and the actual EEG segment) is chosen according to the user’s objectives. Once these parameters are defined, we can choose if FilterBlink will be more or less conservative, respectively losing some VEOG events in the blink detection or detecting other EEG elements with smaller amplitude and correlation with the template. In other words, a threshold of similarity between template and EEG segment (given by L) allows the user to set the sensitivity and specificity of the filter. In this case, the lowest value (*L* = 0.1) was the most effective for the model, although the signals remained stable only up to 0.6.

The non-linear VEOG residues that remain in the ongoing EEG after FilterBlink application limit the application of the present method for ERP analysis in time domain as these residues would yield a persistent low-frequency artifact for the power analysis at the frequency domain in the anterior channels, similar to the original VEOG spectra. It is difficult to say how much this “residue” is not a positive or negative long latency ERP that becomes evident with VEOG averaging (see figure 9). Prior to the VEOG averaged positions, a negative/positive deflection is observed, probably related (at least in part) to the ERPs that occur frequently right before the blinking.

The FilterBlink method is a very intuitive and simple solution whose efficiency has been predicted and the results on the real signals seem also to be suitable, so much so that we believe there is no need to directly compare our method with any other strategy for VEOG suppression.

## Acknowledgements

We wish acknowledge profs. Drs. Antonio Infantosi, Mauricio Cagy and Antonio Mauricio Miranda de Sá for our fruitfull discussions about this method and the EEG model for test it. This work was granded by the “Segundo Programa Interno de Incentivo a Pesquisa” (IFF/FIOCRUZ).

